# Myosin II filament dynamics in actin networks revealed with interferometric scattering microscopy

**DOI:** 10.1101/199778

**Authors:** L. S. Mosby, N. Hundt, G. Young, A. Fineberg, M. Polin, S. Mayor, P. Kukura, D. V. Köster

## Abstract

The plasma membrane and the underlying cytoskeletal cortex constitute active platforms for a variety of cellular processes. Recent work has shown that the remodeling acto-myosin network modifies local membrane organization, but the molecular details are only partly understood due to difficulties with experimentally accessing the relevant time and length scales. Here, we use interferometric scattering (iSCAT) microscopy to investigate a minimal acto-myosin network linked to a supported lipid bilayer membrane. Using the magnitude of the interferometric contrast, which is proportional to molecular mass, and fast acquisition rates, we detect, and image individual membrane attached actin filaments diffusing within the acto-myosin network and follow individual myosin II filament dynamics. We quantify myosin II filament dwell times and processivity as a function of ATP concentration, providing evidence for the predicted ensemble behavior of myosin head domains. Our results show how decreasing ATP concentrations lead to both increasing dwell times of individual myosin II filaments and a global change from a remodeling to a contractile state of the acto-myosin network.

**Statement of Significance:** Here, we show that interferometric scattering microscopy in combination with single particle tracking enables label-free, high contrast imaging of filament dynamics on surfaces, while distinguishing different species based on their mass. These results significantly broaden the available toolkit, and associated capabilities of researchers studying dynamics of biological machines at interfaces.

## Introduction

The dynamics of the cell surface and many cellular processes depend on the interplay between the plasma membrane and the tightly associated dynamic actin cortex (1, 2). Reconstituted, minimal systems of membrane-bound acto-myosin networks are often employed to study physical principles controlling the dynamics of such active composites. Despite their relative simplicity these systems can adopt a range of active states depending on ATP concentration, actin to myosin ratio, actin filament length distribution and concentration (3, 4). As a result, identifying the processes that lead from a remodeling, fluid-like acto-myosin network to a contractile, solid-like network has remained a considerable experimental challenge.

Most experimental and theoretical studies of myosin motor properties have relied on experiments with single-motor head domains. Recent advances in the theoretical understanding of myosin II filaments formed of multiple head domains indicate that small changes in the single-head duty ratio can lead to a switch from non-processive state characterized by weak actin binding no continuous motion along actin filaments to a state of continuous actin binding, efficient motion along actin filaments and force generation due to cooperative effects (5–8). To experimentally test the dynamics and properties of multi-headed myosin II filaments, it is necessary to visualize the network components with a sub-second time-resolution over timescales of tens of minutes, which has been challenging to achieve with fluorescent probes due to photo-bleaching and photo-toxicity.

Here, we employed interferometric scattering (iSCAT) microscopy (9, 10), a label-free imaging technique that makes use of the interference between reflected and scattered light from nanoobjects near an interface. The key advantages of light scattering over fluorescence detection in this context are the lack of an upper limit to the fluorescence emission rate and the absence of photobleaching and thus phototoxicity, enabling long observation times. We quantify microscopic quantities, such as actin filament mobility, myosin filament dwell times and processivity, while simultaneously monitoring mesoscopic phenomena such as network flows and clustering. This approach allows us to link changes in the mechanochemical properties of myosin II filaments to transitions in the acto-myosin network, namely from the remodeling to the contractile state.

## Materials and Methods

### Protein Purification

Actin was purified from chicken breast following the protocol from Spudich and Watt (11) and kept on ice in monomeric form in G-buffer (2 mM Tris Base, 0.2 mM ATP, 0.5 mM TCEP-HCl, 0.04% NaN_3_, 0.1 mM CaCl2, pH 7.0). And myosin II was purified from chicken breast following a modified protocol from Pollard (12) and kept in monomeric form in myo-buffer (500 mM KCl, 1 mM EDTA, 1 mM DTT, 10 mM HEPES, pH 7.0). The day prior to experiments, functional myosin II proteins were separated from proteins containing dead head domains by a round of binding and unbinding to F-actin at a 5:1 actin to myosin ratio (switch from no ATP to 3 mM ATP) followed by a spin at 60,000 rpm for 10 min at 4 °C in a TLA100.3 rotor. The supernatant containing functional myosin II is dialyzed against myo-buffer overnight and used for experiments for up to three days.

To control the length of actin filaments, we titrated purified murine capping protein to the actin polymerization mix as described in (4). To link actin to the SLB, we used a construct containing 10x His domains followed by a linker (KCK) and the actin binding domain of Ezrin (HKE) as described earlier (13).

### Supported Lipid Bilayer and Experimental Chamber Preparation

Glass coverslips (#1.5 borosilicate, Menzel, Germany) for SLB formation were cleaned with Hellmanex III (Hellma Analytics, Mühlheim, Germany) following the manufacturer’s instructions followed by thorough rinses with EtOH and MilliQ water, blow dried with N_2_ and finally passed briefly over a Bunsen burner flame. For the experimental chamber, 0.2 ml PCR tubes (Tarsons Products, Kolkata, India) were cut to remove the lid and conical bottom part. The remaining ring was stuck to the cleaned glass using UV glue (NOA88, Norland Products, Cranbury, NJ) and three minutes curing by intense UV light at 365 nm (PSD-UV8T, Novascan, Ames, IA). Freshly cleaned and assembled chambers were directly used for experiments.

Supported lipid bilayers (SLB) containing 98% DOPC and 2% DGS-NTA(Ni^2+^) lipids were formed by fusion of small uni-lamellar vesicles (SUV) as described previously (4). Prior to experiments with F-actin we formed SLBs in chambers filled with 100 μl KMEH (50 mM KCl, 2 mM MgCl_2_, 1 mM EGTA, 20 mM HEPES, pH 7.2). SLB formation was observed live using ISCAT microscopy, to ensure that sufficient SUVs fused to form a uniform, continuous and fluid lipid bilayer (Video 1).

### Formation of Acto-Myosin Network

In a typical experiment, SLBs were formed, incubated with 10 nM HKE for 40 min and washed three times with KMEH. During this incubation time, F-actin was polymerized. First 10%_vol_ of 10x ME buffer (100 mM MgCl_2_, 20 mM EGTA, pH 7.2) were mixed with the G-actin and, optionally, together with the capping protein stock and incubated for 2 min to replace G-actin bound Ca^2+^ ions with Mg^2+^. Addition of 2x KMEH buffer supplemented with 2 mM Mg-ATP induced F-actin polymerization at a final G-actin concentration of 5 μM. After 20-30 min incubation, the desired amount of F-actin was added to the SLBs using blunt-cut 200 μl pipette tips. An incubation of 30 min allowed the F-actin layer to reach an equilibrium state of binding to the SLB. For the addition of myosin II filaments to the sample, intermediate dilutions of myosin II proteins were prepared in milliQ water at 10 times the final concentration from fresh myosin II stock (C_myoII_ = 4 mM; 500mM KCl, 1mM EDTA, 1 mM DTT, 10 mM HEPES, pH 7.0) in a test tube and incubated for 5 min to enable myosin II filament formation. Then, 1/10 of the sample buffer was replaced with the myosin II filament solution and supplemented with Mg-ATP (100 mM) at 0.1 mM final concentration. To summarize, the final composition of the assay is 50mM KCl, 2mM MgCl_2_, ImM EGTA, 20mM HEPES, 0.1mM ATP at pH 7.2 with F-actin (C_G-actin_ = 100-300nM) and myosin II filaments (C_myoII_ = 0-100nM). Afterwards, the dynamics of the acto-myosin system were observed for up to 60 min. The system usually showed a remodeling behavior for the first 10-15 min before contraction and aster formation (due to ATP concentrations dropping below 10 μM as estimated from the activity of myosin II and earlier reports (14)). Once, the system reached a static, jammed state and no myosin activity could be observed, the system could be reset into a remodeling state by addition of Mg-ATP (100 mM) to a final concentration of 0.1 mM (Video 10). Each step of this procedure was performed on the microscope, which allowed us to check the state continuously. The open chamber design allowed the addition of each component from the top without induction of flows that would perturb the actin network. Evaporation was below 5% during a period of 60 minutes, so that salt and protein concentrations can be considered constant over the time course of a typical experiment (Fig. S1A). All experiments were performed at room temperature (22° C).

Details about the actin, myosin and ATP concentrations used in each experiment can be found in table (S1).

### iSCAT microscopes

The principle of interferometric scattering microscopy is based on the interference of light reflected from the glass substrate with the light that is scattered from the particle of interest, and the measured intensity at the detector is given by

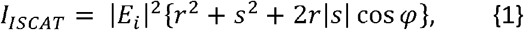

Where |E_i_|^2^*r*^2^ describes the light intensity reflected by the glass-water interface (r ~ 0.065), |*E_i_*|^2^*s*^2^ is the pure scattering contribution (neglectable for very weak scatterers) and 2*r*|*s*| sin *φ* denotes the interference between scattered and reflected light with *φ* being the phase difference between the two (15). iSCAT experiments were performed on two different home-built setups similar to those detailed in (16). Briefly, a weakly focused laser beam was scanned across the sample over an area of 24 x 24 μm^2^ (445 nm laser) or 32.6 x 32.6 μm^2^ (635 nm laser). The light reflected from the glass-water interface together with the scattered light from the sample was imaged onto a CMOS camera (445 nm laser: Point Grey GS3-U3-23S6M-C, Flir, Canada; 635 nm laser: MV-D1024-160-CL-8, Photonfocus, Switzerland). The cameras were controlled using home-written LabVIEW software. The setup with the 445 nm laser had a 3.5 mm partially reflective mirror placed in the re-imaged back focal plane of the objective for enhanced scattering contrast as described in (16). The videos were recorded at 50 fps (445 nm laser) and 25 fps (635 nm laser), and pre-averaged by a factor of 5 to reduce noise. The illumination intensity on the sample (445 nm laser: 250 W/cm^2^; 635 nm laser: 1.9 kW/cm^2^) was set to nearly saturate the camera with the returning light. The pixel sizes were 23.4 nm/pixel (445 nm laser) and 31.8 nm/pixel (635 nm laser).

### Image processing

To reduce noise, five images were averaged per time point, i.e. for an effective frame rate of 10 Hz images were recorded at 50 Hz. Non-sample specific illumination inhomogeneities, fixed-pattern noise and constant background were removed from the raw images by dividing each of them with a flat field image that contained only these features. The flat field image was computed by recording 2000 frames of the sample while moving the stage. For each pixel, a temporal median was calculated resulting in a flat field image that only contained static features.

### Median filtering

Movies were median filtered using MATLAB (MathWorks, Natick, MA, USA). For each image sequence, the median is computed for each pixel, subtracted from the original image sequence and the median filtered image sequence as well as the computed median filter are saved.

### Actin filament tracking

Actin filaments that became visible after median filtering and that did not cross other actin filaments for at least 1000 frames were tracked using image J (http://imagej.nih.gov) and the plugin JFilament (17, 18). The obtained tracking traces were analyzed using MATLAB to compute the position of the center of mass (CM) and the filament orientation for each time point and to generate plots of the CMs total mean-squared displacement as well as the parallel and perpendicular components of the mean-squared displacement with respect to the filament orientation.

### Actin filament length measurements

Image stacks of actin filaments landing on the HKE decorated SLBs were taken at 10 Hz immediately after addition of actin filaments to the sample. Theses image stacks were split in segments of 10 s and the median of each segment was subtracted from its last frame, to visualize freshly landed, isolated actin filaments. The images were then converted from the 32-bit interferometric contrast values to 8-bit (by the formula *f(x) = 1000*(-x) + 1000)*, bandpass filtered (low pass: 3 pixel, high pass: 20 pixel) and analyzed with the image J plugin Neuron J (19).

### Actin filament layer thickness measurements

The maximum interferometric contrast values of randomly drawn line scans across SLB bound actin networks before the addition of myosin were taken and divided by the average interferometric contrast value of a single actin filament (see Fig 1D) to obtain an estimate of the actin network thickness as number of actin filaments. It is to note that the obtained maximum thickness of 6 actin filaments would amount to a network height of about 6 × 8 nm = 48 nm, which lies within the working distance of the objective used.

**Figure 1.**
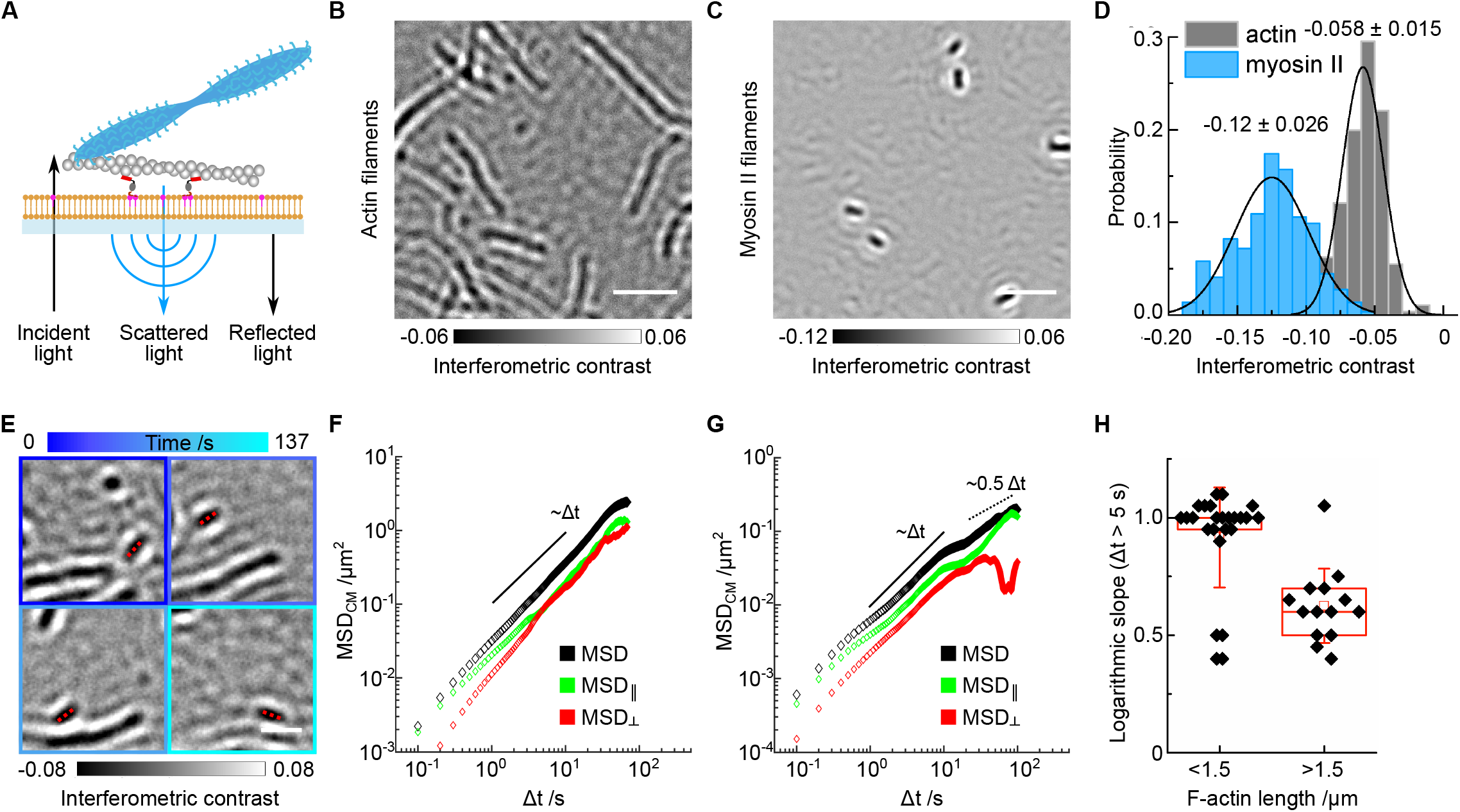
experimental setup: **A)** Diagram of the *in-vitro* system consisting of a supported lipid bilayer (orange), actin-membrane linker protein deca-histidine – KCK - Ezrin actin binding domain (HKE; dark gray and red), actin filaments (gray) and muscle myosin II filaments (blue), arrows indicate principle of iSCAT microscopy. **B-C)** Example images of actin filaments (B) and myosin II filaments (C), both recorded with a 445 nm laser iSCAT system; scale bar: 2 μm. **D)** Histogram depicting the interferometric contrast distribution along actin filaments (gray, N_measure_ = 562 measurements along N_fil_ = 12 filaments) and myosin II filaments (blue, N_measure_ = 303 measurements along N_fil_ = 14 filaments). **E)** Example of tracking a single actin filament (red dashed line, l = 0.4 μm) inside an actin network imaged over 137 s (from dark blue to cyan: 0-137 s); scale bar: 1 μm. **F)** Corresponding mean-squared displacement (MSD) of the filament’s center of mass (black) and the components parallel (green) and perpendicular (red) to the actin filament orientation. **G)** Mean-squared displacement of the center of mass of a long actin filament (l= 4 μm, Video 4) (black) and its components parallel (green) and perpendicular (red) to the actin filament orientation, indicating confinement at timescales >10 s. **H)** Box plot comparing the diffusive behavior (characterized by logarithmic slopes of MSD plots) of short (<1.5 um, N = 25) and long (> 1.5 um, N = 14) actin filaments indicating confined diffusion for filaments longer than 1.5 μm due to the surrounding actin meshwork.

### Myosin II filament length measurements

Line-scans along the long axis of single myosin filaments were taken and the distance between the half maximum points at both ends were taken as the length of the myosin II filament.

### Myosin binding dynamics

Imaging with an effective frame rate of 5-10 Hz for several minutes was sufficient to capture a broad range of myosin filament dynamics with high accuracy. To remove any signal originating from static structures such as immobile actin filaments or small impurities in the SLB, the image sequences were median filtered (20). In a second step, a maximum projection of the time series was used to visualize the tracks occupied by myosin II filaments during the experiment. Lines following these tracks were then used to compute kymographs depicting the myosin II filament binding times and their motion along actin filaments (kymograph tool in Image J, line width 3). Dwell time, run length and velocity distributions were plotted and further analyzed using OriginPro 2018 (OriginLab Corporation, Northampton, MA, USA) and MATLAB. Best fitting functions for the myosin II filament dwell times were selected using MEMLET, which estimates the maximum likelihood of a fitting function to describe a data point distribution (21).

### Detection and Tracking

Image analysis and single particle tracking (SPT) was carried out using ImageJ and Python code developed for this work (detailed description in (22), code is available upon request). Briefly, individual myosin-filament detection was implemented in the Python programming language using the Sep package (based on the algorithms of Source Extractor)(23–25), which generates the position, spatial extent, and orientation of the ellipsoidal myosin II filaments for each frame. Based on the amplitude of spatial displacement, oritentation changes and the detected particle area tracks were generated and classified in regions of random or directed motion.

## Results

### Detection and characterization of actin and myosin filaments

The critical step to use label-free imaging for acto-myosin dynamics on a membrane requires detection and distinction of actin and myosin II filaments. Due to the absence of crowding factors or excess proteins in our experiments, the actin and myosin II filaments were the principal sources of light scattering after subtraction of background signatures originating from cover glass roughness leading to interference with the light reflected from the cover glass surface and referred to in the following as interferometric contrast with values <0 (Fig. 1A, Fig. S1B)(20, 26). Myosin II and actin filaments landing on bare glass slides exhibited interferometric contrasts of −0.12 ± 0.0263 and −0.058 ± 0.015, respectively, using an iSCAT setup equipped with a 445nm laser (Fig. 1B-D). The interferometric contrast values were smaller using a 635nm laser setup with values of −0.017 ± 0.006 for myosin II filaments and −0.002 ± 0.001 for actin filaments, respectively (Fig. S1C). The contrast ratio (^0.017^/_0.002_ = 8.5) was larger than with the 445nm laser setup and reflected the mass ratio of actin to myosin II filaments per unit length (within a diffraction limited spot of 210 (± 10) nm: 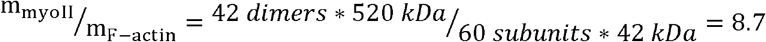) (27–29) and the linear scaling of the interferometric signal with molecular mass (26). The weaker contrast difference measured with the 445nm laser (^0.12^/_0.06_ = 2) was likely due to a non-negligible contribution from direct scattering by the large myosin II filaments (the |s|^2^ term in equation {1}) leading to a reduced interferometric contrast at high molecular masses. In our buffer conditions, containing 50 mM KCl, myosin II filaments exhibited an average length of 520 ± 130 nm (N=269) (Fig. S1D) as reported earlier (3). The clear difference in interferometric contrast with the 635 nm laser setup and the characteristic, uniform shape of myosin II filaments provided a solid basis to distinguish myosin II filaments reliably from actin filaments, while the 445nm laser setup was mainly used to track actin filament dynamics.

Next, we characterized the formation of actin networks on supported lipid bilayers (SLB). After directly observing successful SLB formation (Video 1), incubation with an actin-membrane linking protein (deca-histidine – KCK – Ezrin actin binding domain, HKE) and addition of actin filaments, we did not detect any measurable effect of the SLB on the interferometric contrast of actin filaments, and regions exhibiting consecutive deposition of multiple actin filaments within a diffraction-limited spot displayed a step-wise increase in the interferometric signal (Fig. S1E, F; Video 2). By dividing local contrast measurements by the average value for a single actin filament (−0.058 ± 0.015), we estimated that, under the conditions used here ([G-actin] = 100-350 nM), the actin layers were 1 to 6 filaments thick, with 2 filaments per diffraction-limited spot being most frequent (Fig. S1G). This is entirely consistent with previous results using fluorescently labelled actin filaments (4).

It was important to check for the fluidity of the supported lipid bilayers to ensure the lateral mobility of the membrane-actin linker HKE and, hence, the remodeling of the membrane tethered actin network. Since HKE could not be visualized directly using iSCAT due to its low molecular weight (15kDa) (13), we assessed the mobility of individual short actin filaments as a proxy. Offline median subtraction (median of 2 min image sequences) removed quasi-static actin network components and revealed the mobile filament fraction (Fig. 1E, Video 3). Tracking of individual filaments and analysis of their mean-squared displacement indicated a fluid lipid bilayer with free diffusive behavior for most actin filaments shorter than 1.5 μm on timescales of 5 s and longer reaching usually MSD values of > 0.1 μm^2^ (Fig. 1E, F, H; S1H, J). Longer filaments also displayed diffusive behavior on short timescales but were more confined with lower MSD values (in the range of 0.1 μm^2^ at Δt = 5s) characterized by lower slopes of the MSD (Fig. 1G, H, S1I, K; Video 4). Two major effects are likely to influence actin filament mobility here, the number of molecules tethering the actin filament to the membrane and steric effects by the surrounding actin network. Considering that longer actin filaments will have a higher number of HKE molecules tethering them to the bilayer, one can assume that the actin filament motility would decrease as a function of actin filament length as was discussed in the case of microtubules in (30). The presence of a heterogenous meshwork resulting in different confinements for actin filaments depending on their local environment could give rise to tracks showing non-Gaussian diffusion as described in (31) and as we observe (Fig. S1 J, K). However, a detailed analysis of whether actin filament diffusion could be used to reveal the actin network mesh size distribution or how actin mobility depends on actin filament length would be beyond the scope of this study.

### Binding dynamics of myosin II filaments to actin

We then characterized myosin II filament binding to membrane-bound actin networks at 100 μM ATP (t= 1 min), which fueled continuous network remodeling for several minutes until it became contractile at about t = 16 min, due to low ATP levels (Fig. S2A, B; Video 5). Previously, we analyzed this transition e.g. by calculating the spatial density correlation of actin which changed clearly at the onset of the contractile state due to the clustering of actin (4). Given that we observed in the present work similar network dynamics and time scales of the remodeling and contractile states, we did not perform a similar analysis, but wanted to understand, whether changes in myosin II filament dynamics could be associated to changing network dynamics. We followed myosin II filament dynamics using a 635 nm laser iSCAT setup over a time window of >16 min by recording multiple sequences of 2 min movies (3000 frames at 25Hz) due to computer hardware related limitations for rapid data storage. After averaging over 5 frames to reduce noise levels, we obtained an effective frame rate of 5Hz, which was 10-20 times faster compared to earlier studies using fluorescence light microscopy limited by photo-toxicity (4, 32). As described previously (4, 14, 33), we defined the onset of network contractility by the myosin induced formation of actin clusters that continued to merge into larger structures until the system reaches a jammed or static state without any visible actin or myosin motion. By contrast, the remodeling state was characterized by local, transient changes of the actin network without inducing any large-scale contractile flows. Changes in network contractility were due to changes in the available ATP, and addition of fresh ATP could reverse the strong binding of myosin II filaments (Video 6).

By creating kymographs along the tracks of myosin II filaments (Fig. S2C, D; Video 7), we found that the distribution of myosin II dwell times on F-actin followed a double-exponential decay function. At t = 1 min, the computed time constants were τ_off1_ = 1.23 s ([1.12 s - 1.36 s]_95%_) and τ_off2_ = 12.6 s ([11.3 s - 14 s]_95%_) with 66% - 71% of the events being described by τ_off1_ (Fig. 2A). At t = 16 min, the myosin II filament dwell time constants were τ_off1_ = 3.7 s ([3.1 s - 4.2 s]_95%_) and τ_off2_ = 11.9 s ([9.4 s - 15.5 s]_95%_) with 63 % - 87 % of the events being described by τ_off1_ (Fig. 2B).

**Figure 2.**
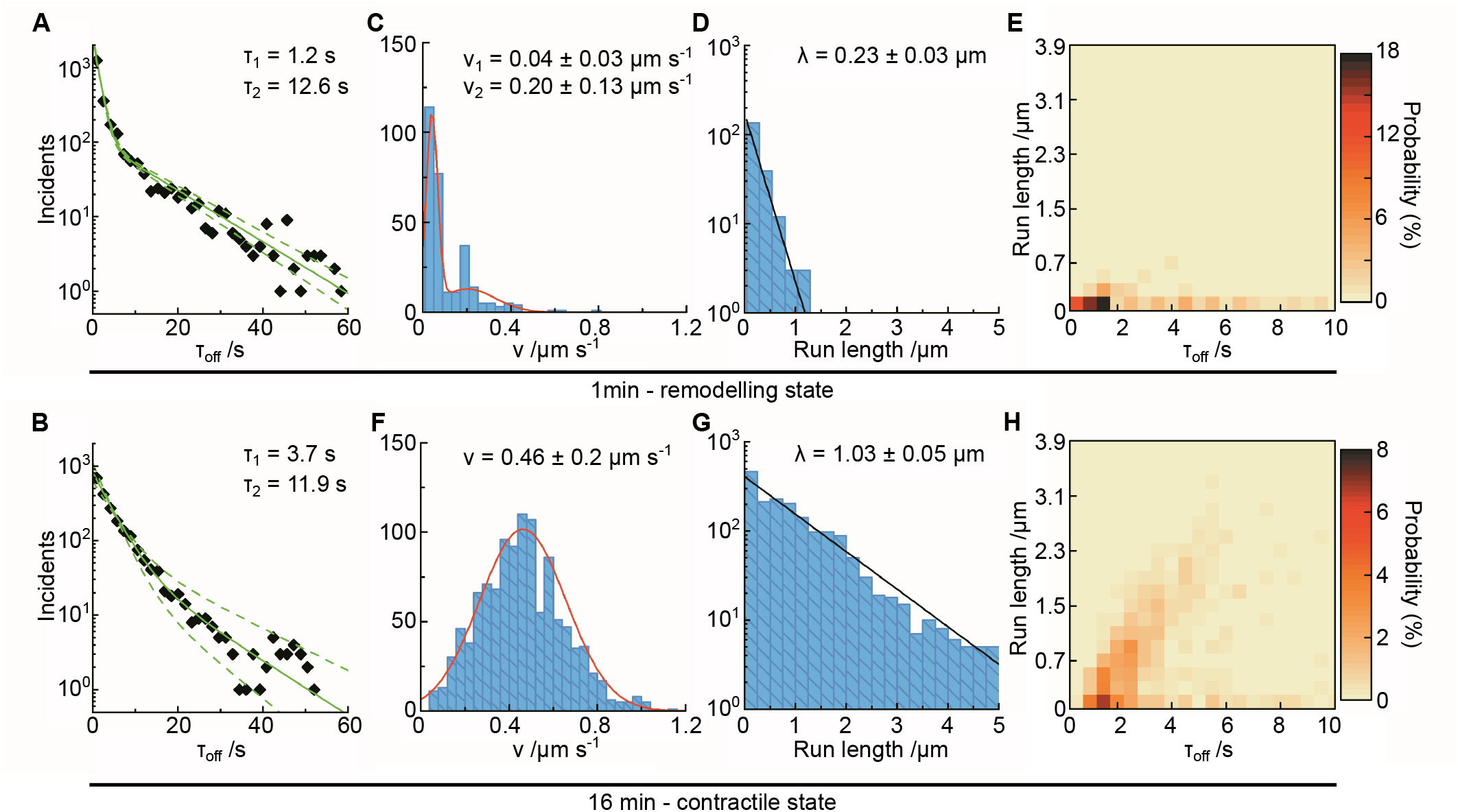
myosin II filament dynamics in the acto-myosin network at different ATP concentrations: **A, B)** Histogram of myosin II filament dwell times on actin at t = 1min (100 μM ATP) (N = 6400) **(A)** and at t = 16 min (N = 8000) **(B);** each diamond represents a 5 frame = 1 s bin. **C-E)** Histograms of the myosin II filament velocities **(C),** run lengths **(D)** and frequency plot of run length vs dwell time **(E)** extracted from a subset of the myosin **II** filament kymographs at t = 1min (N=432). **F-H)** Histograms of the myosin II filament velocities **(F),** run lengths **(G)** and frequency plot of run length vs. dwell time **(H)** extracted from a subset of the myosin II filament kymographs at t = 16 min (N=1133). Best fits for the distributions were computed with the MEMLET fitting routine: (A and B), double exponential decay (solid green line, dashed lines indicate 95% confidence level), (C) double Gaussian (red line), (F) single Gaussian (red line) and (D and G) single exponential decay (black line). Data displayed in (E and H) is displayed in the range of 0-10 s in order to highlight this dwell time regime.

We went on to analyze the motion of myosin II filaments. At t = 1min, the mobile myosin II filaments exhibited a velocity distribution that can be described by a sum of two Gaussians with >60 % of myosin II filaments travelling at v_myoII,1_ = 0.04 (±0.03) μm s^-1^ and the remaining at v_myoII,2_ = 0.20 (±0.13) μm s^-1^ (Fig. 2C). The corresponding run length distribution decays exponentially with a characteristic run length of λ = 0.23 (±0.03) μm (Fig. 2D). Plotting run length versus dwell time (i.e. total time of attachment during which the run happened) shows a moderate positive correlation (Pearson coefficient 0.38, p-value: 7*10^-10^) for dwell times < 3 s and a weak negative correlation (Pearson coefficient −0.17, p-value: 2*10^-2^) for longer binding times (Fig. 2E, Fig. S2E). This implies that the population corresponding to short dwell times is mobile while the population with long dwell times exhibits slow, reduced motion and eventually becomes immobilized.

At t = 16 min, mobile myosin II filaments travelled with an average velocity of v_myoII_ = 0.46 (±0.20) μm s^-1^ (Fig. 2F, single Gaussian distribution) and the characteristic run length was *λ* = 1.03 (±0.05) μm (Fig. 2G). The correlation between run length and dwell time is stronger compared to t = 1 min with most myosin II filaments exhibiting persistent motion for up to 6 s (Pearson coefficient 0.57, p-value: 1*10^-16^) and a loss of this correlation at longer dwell times (Pearson coefficient −0.18, p-value: 3*10^-3^) (Fig. 2H, Fig. S2F).

To assess whether the measured velocities were a combination of myosin and actin network motion, we applied a 50 frames median filter (corresponding to 10s) filtering out the fast-moving myosin II filaments and highlighting actin filament motion. The largest actin network displacements that we could observe occurred only locally and sporadically and did not exceed 50 nm s^-1^ in the contractile state (t = 16 min) and 15 nm s^-1^ in the remodeling state at t = 1 min (Fig. S2G). These observations agree with earlier studies (4, 34) and underline that myosin II filament velocities result directly from motor activity on actin filaments. A comparison of the forces required to drag actin filaments and the attached actin-membrane binding proteins along the supported lipid bilayer with the drag forces experienced by myosin filaments moving through an aqueous buffer suggest as well, that myosin activity will propel mainly myosin II filament and will not be considerable reduced by slippage of the membrane tethered actin filaments (see supplementary information) (30).

### Myosin II filament tracking reveals links between binding mode, orientation and mobility

Even though the use of kymographs to analyze binding dynamics and the mobility of motor proteins is well established, it bears some drawbacks particularly when studying dynamics in complex, remodeling networks: Kymographs only work well along static tracks making it difficult to capture movements along actin filaments that change position over time; in addition, short binding events outside the kymograph lines would not be detected or, if captured, the reduced dimension due to the line scan would make it difficult to distinguish from noise. The characteristic shape and signal of myosin II filaments obtained with iSCAT microscopy, however, allowed us to develop an automated single-particle tracking (SPT) algorithm to analyze the dynamics and orientation of individual myosin II filaments in a more detailed way than possible with kymographs. By analyzing the intensity distribution of pixels belonging to each myosin II filament, we could extract the particle location and orientation for each time point. Tracks were generated based on criteria including the particle size and the maximum displacement between frames) (Fig. 3A, B; Video 8) (22). Given the chosen intensity and area thresholds, detection of unbound myosin II filaments moving by chance near the glass surface was very unlikely, as it would move out of a 200nm radius within less than 10ms, which was significantly shorter than our image integration time (see supplementary information). By quantifying the diffusivity of particles exhibiting random motion, individual myosin II filament tracks could be divided into periods of random and directed motion (Fig. 3B, S3A-C). We used the average MSD data at t = 1 min (100 μM ATP) to calculate a threshold for identifying directed motion to classify individual tracks. The random motion or weak-binding state was characterized by an increased rate of large angular fluctuations, and random displacements that resulted in linear time evolution of the MSD, resembling a diffusive particle.

**Figure 3.**
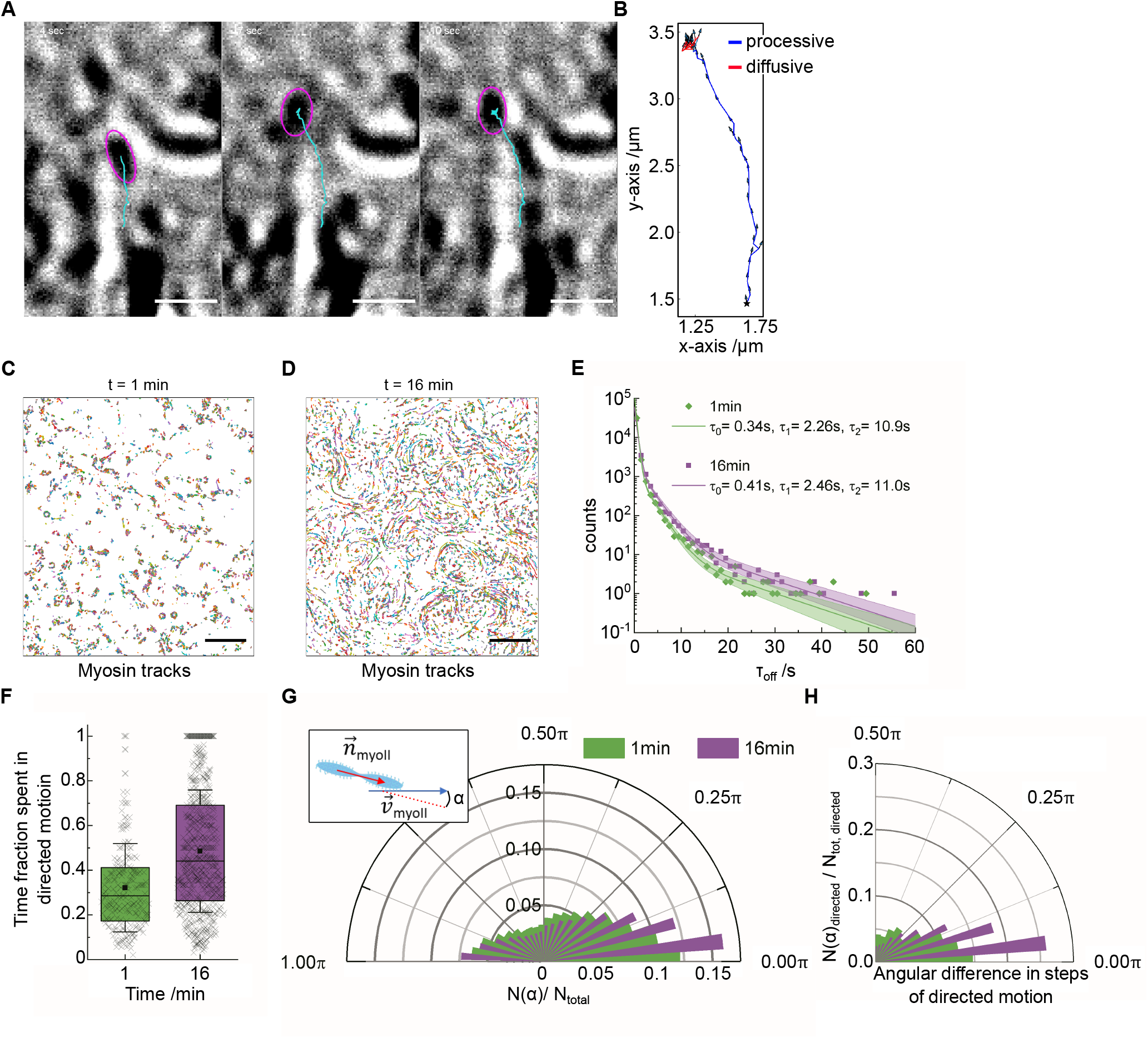
single particle tracking of myosin II filaments provides further insights into change of myosin dynamics with decreasing ATP concentration: **A)** Example image sequence showing the tracking of a myosin II filament (purple circle locates the current detection, cyan line depicts the particle’s traces), scale bar: 1 μm. **B)** Corresponding track with segmentation into segments of directed (blue) and random (red) motion. **C, D)** Images depicting all tracked myosin II filaments (in alternating colors to distinguish individual tracks) at **(C)** t = 1 min (100 μM ATP) (N = 14016) and **(D)** t = 16 min (N = 15647), scale bar: 5 μm. **E)** Corresponding histogram of myosin II filament dwell times on actin (t = 1 min: green; t = 16 min: purple); best fits for the distributions were computed with the MEMLET fitting routine: triple exponential decay with t_min_ = 0.3s (solid line), boundaries depict 95% confidence level. **F)** Box plot depicting the ratio of time of directed motion versus dwell time for myosin II filaments displaying directed motion (N(1min) = 221; N(16min) = 640). **G)** Radial histogram depicting the angular difference α between myosin II filament orientation and its velocity vector for all detected myosin II filament steps. **H)** Radial histogram plot depicting the ratio of steps in directed motion versus all detected steps for each angular difference α, indicating that there is a clear increase of filaments that are aligned along their axis of propagation in the contractile state.

This automated detection method provided a more comprehensive picture of myosin II filament dynamics in the remodeling state at t = 1 min and at the contractile state (t = 16 min) (Fig. 3C, D). Moreover, it increased the sensitivity for short binding times compared to the kymograph method. This higher detection sensitivity was reflected in the number of myosin II filaments identified in each frame, which fluctuated between 94 (±14) and 214 (±30) detections between t = 1 min and t = 9 min and rose to 392 (± 29) detections per frame at t =16 min (Fig. S3D). At t = 1 min (100 μM ATP), we found characteristic binding times of τ_off0_ = 0.34 s (0.33 s – 0.35 s), τ_off1_ = 2.26 s (2.13 s - 2.41 s) and τ_off2_ = 10.9 s (9 s - 12.8 s) and at t = 16 min τ_off0_ = 0.41 s (0.39 s - 0.43 s), τ_off1_ = 2.46 s (2.33 s - 2.63 s) and toff2 = 11 s (10 s – 12.5 s) (Fig. 3E). Interestingly, the characteristic times only increased slightly at t = 16 min, but significantly (performing a non-parametric test (Kolmogorov-Smirnov) shows that the t = 16 min dwell time distribution is larger than the t = 1 min distribution with p = 0.04 for dwell times > 2 s). The number of events with binding times longer than 3 s increased > 1.6-fold compared to the remodeling state (Fig. S3E). Even though the number of long duration binding events in the contractile state could be an underestimate, as only binding-unbinding cycles occurring within the recording time were considered, discarding myosin II filaments that were bound for the entire length of the movie, that were traveling into clusters or out of the field of view.

The fraction of myosin II filaments moving in a directed manner, however, increased nearly 3-fold (N_directed_/N = 221/35644 at t = 1 min; N_directed_/N = 640/35358 at t = 16 min). Concomitantly, the fraction of time each myosin II filament performed directed motion increased from 0.3 (±0.2) at t = 1 min to 0.5 (±0.3) at t = 16 min (Fig. 3F, S3F) of the total bound time, while the average number of switches between the two states of motion changed only from 1.1(±0.3) (N=221) to 1.2(±0.5) (N=640) suggesting that the additional time of directed motion at t = 16 min is spent in longer, continuous runs and not split up in multiple short sequences of directed and undirected motion (Fig. S3G). This analysis is limited to runs lasting ≥ 10 frames (2 s)(22) and experiments at higher frame rates would be necessary to reveal dynamics at shorter timescales.

These results indicate that the contractile state is driven by both an increased total number of myosin II filaments showing directed motion and a higher persistence of each myosin II filament. The observed changes of myosin II filament run length and dwell time are most likely due to changes in ATP concentration and not due to changes in the actin filament length as the characteristic actin filament length at the onset of the experiments was 7 (± 3.5) μm, and hence exceeded the run lengths we measured for myosin II filaments at t = 16 min (Fig. 2G). To assess, whether the properties of individual myosin motors changed during the course of the experiment, we analyzed the myosin II filament displacement per frame of all tracks during periods of directed motion and did not find any detectable changes over the course of the experiment (Fig. S3H). This implies that the velocity distribution of myosin II filaments at directed motion remained constant over the range of 10-100 μM ATP and indicates that the difference in velocities measured with the kymograph method (Fig. 2 C, F) might have been due to tracks comprising intervals of directed and random motion.

By computing the angle α between myosin II filament orientation and its velocity direction for each myosin II filament step, we found that α varied significantly (± 45° FWHM) at t = 1 min, whereas it narrowed with increasing time to ±25° (FWHM) at t = 16 min (Fig. 3G, S3I). Importantly, at t = 16 min, the majority of myosin II filaments moving in a directed fashion are aligned with their direction of propagation (Fig. 3H, S3J). This improved alignment with the actin tracks at t = 16 min could be due to higher numbers of myosin head domains binding to actin filaments, which would support long range transport and buildup of persistent material flows. As an approximation, we computed the effective rotational flexibility of myosin II filaments if the observed angular fluctuations (α) would solely be driven by thermal energy (*k_B_T*) using the relation *k_B_T* = 〈Δ*α*^2^〉*K_tor_* with K_tor_ being the effective torsional spring constant of the myosin II filament around its attachment point and found that *K_tor_* increased from 7.5 to 9.2 *pN/nm rad*^2^ during the time of the experiment (Fig. S3K). This is about 3 times below the reported value for an individual myosin II head domain of 23 *pN/nm rad*^2^ (35–37) which might be due to several factors: i) experiments performed in solution in contrast to the previous studies using myosin II head domains adsorbed and fixed on EM grids, ii) myosin II filaments containing differently oriented myosin head domains which could ease re-orientation by changing the myosin head binding to the actin filament. However, the observed increase of *K_tor_* with decreasing ATP concentrations could be due to the binding of multiple myosin heads simultaneously to an actin filament. This rotational flexibility is likely to allow myosin filaments to connect multiple actin filaments, which would be important to generate network contraction (38, 39).

While we were following the network dynamics over the course of 20 min, it was a striking to observe that the transition from the remodeling to the contractile state happened very suddenly at around 16 min (Video 5). This was reflected by a sudden change in the ratio of myosin II filaments showing directed motion (Fig. S3F) as well as in the change in filament orientation (Fig. S3J) at that time point. This indicates that gradual changes in the ensemble of myosin head domains (e.g. binding time) lead to a sharp transition between regimes where myosin II filaments either mediate a remodeling or contractile behavior of the network.

### Myosin II filament flows generate transiently stable contractile zones

In acto-myosin networks that reached their contracted state, we observed the formation of contractile zones, into which myosin II filaments moved continuously from multiple directions. These contractile zones were characterized by a cluster of myosin II filaments in their center holding multiple actin filaments together. Interestingly, despite the steady flow of myosin II filaments into the clusters, they did not seem to grow significantly in size or mass. After subtracting the median image of the image sequence showing a cluster region, it became evident that myosin II filaments detached once they reached the core of the cluster preventing overall growth of the myosin II filament cluster (Fig. 4A, B; Video 9). This could be due to the lack of free actin filaments and mechanical tension inside the clusters forcing excess myosin filaments to leave the detection zone either by detaching from the cluster or by stacking up to form three dimensional structures above the membrane tethered actin network as we observed earlier (4, 40).

**Figure 4.**
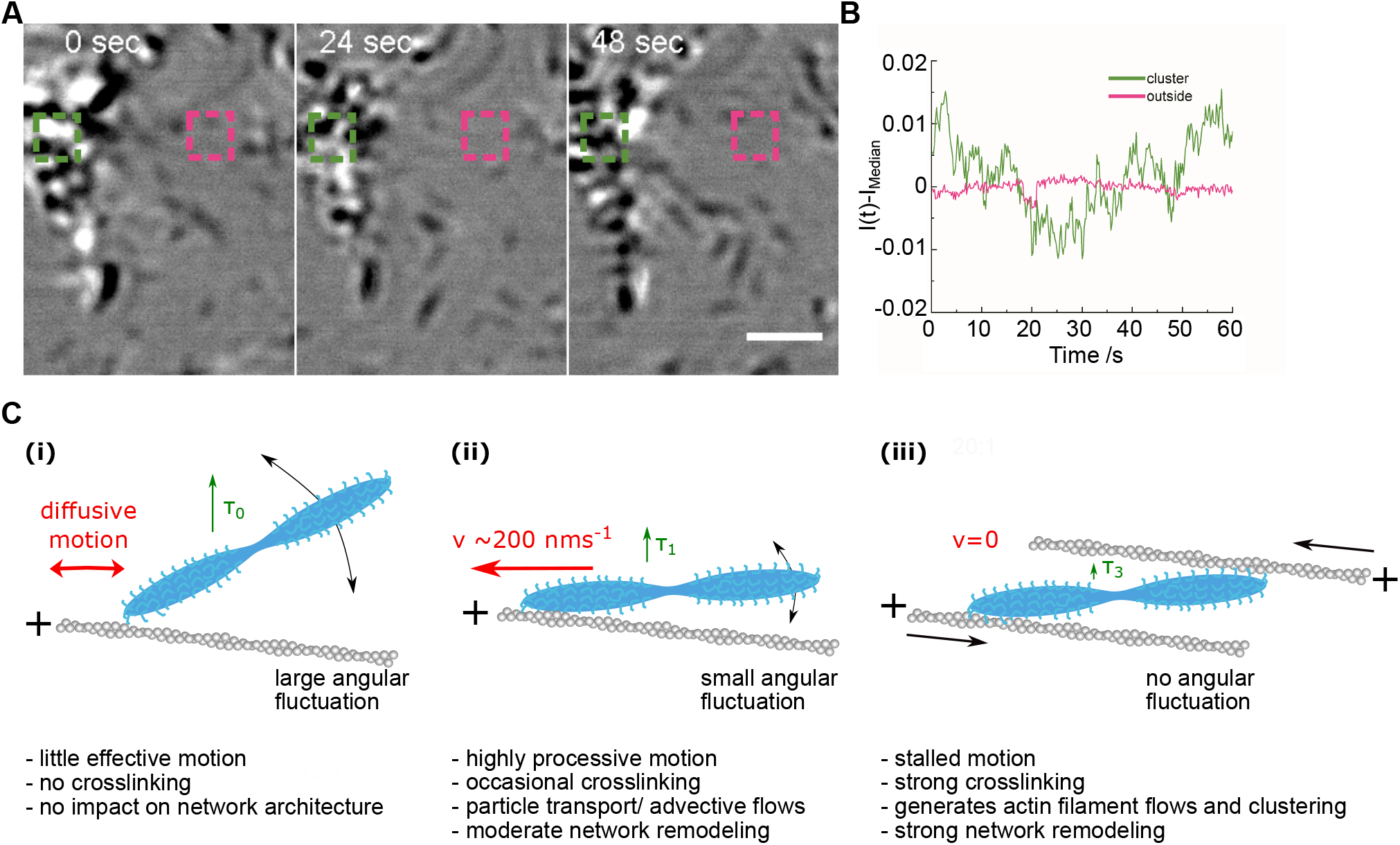
dynamics within acto-myosin clusters: **A)** Image sequence depicting the dynamics of myosin II filaments within an acto-myosin cluster; to make the dynamics in the cluster visible, the time median of the image sequence was subtracted from each frame; 635nm iSCAT setup, scale bar 2 μm. **B)** Graph depicting the change of interferometric contrast from the time median of the regions depicted in **(A)** inside (green) and outside (magenta) the cluster. **C)** Schematic summarizing the different observed binding modes of myosin II filaments and their assumed effect on the acto-myosin network dynamics and organization.

## Discussion

Studying muscle myosin II filament dynamics inside a membrane-bound actin network with iSCAT microscopy has the potential to provide insights on the relation between mesoscale network dynamics and minute changes of the physico-chemical properties of motor head domains in myosin II filaments. In this work, we demonstrated that changes in muscle myosin II filament dwell times and myosin filament motion can be reliably tracked within actin networks and that changes in myosin filament dynamics can be related to changes in the acto-myosin network from a remodeling to a contractile state. This is emphasized by the observation that muscle myosin II filaments move longer and more directed in the contractile state (here at t = 16min) than in the remodeling state at 100 μM ATP. These changes in muscle myosin II filament dynamics are likely to be due to a decrease in ATP concentrations over time as previously reported (14, 33). Recent theoretical studies suggested that factors like changes in ATP concentration and mechanical load, e.g. transmitted via actin filaments, increase the processivity of ensembles of myosin II head domains and can influence myosin II filament binding dynamics (8, 41). Our observations are in line with these predictions, however, the difference in ensemble size described in the theoretical models (8-48 myosin head domains) and studied here (100-250 head domains) poses limits on the comparison. Further studies with nonmuscle myosin II filaments, which form better defined filaments of ~30 dimeric subunits would be needed to elucidate the mechano-sensitive properties of myosin head domain ensembles and test the theoretical predictions which mainly relied on data obtained from single myosin head domain studies (8, 41). The approach introduced here could be used to test these models with data directly obtained from myosin II filaments.

As a proof of concept, we used here muscle myosin II filaments as their size made them easily detectable in iSCAT microscopy. These should be replaced by non-muscle myosin IIA and myosin IIB filaments for more physiologically relevant studies of myosin driven cell cortex dynamics in future (42–44). Detection with iSCAT would be possible as the typical size of a non-muscle myosin II A (NMIIA) filament (~300nm long, 29 dimers @ 490 kDa) would lead to a contrast ratio with an actin-filament within a diffraction limited spot of 210 (± 10) nm of 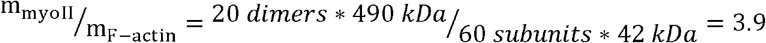
 and the optical resolution of the 445nm laser iSCAT setup would allow to detect NMIIA filaments as elongated structures.

Given the large number of myosin head domains within a single myosin II filament, we would expect complex binding behavior with multiple characteristic attachment times. Our data, however, shows that myosin II filament binding is determined by 3 major time scales: τ_off0_ in the range of 0.3-0.4 s, τ_off1_ in the 2-3 s range and τ_off2_ in the 10 s range. The shortest time scale likely corresponds to the binding - unbinding of a single myosin head domain (45). Considering the dumbbell structure of myosin II filaments, we attribute τ_off1_ to the binding of several myosin head domains at one side of the dumbbell, whereas τ_off2_ would represent the simultaneous binding of myosin heads from both sides. This model would explain the improved alignment of myosin filaments at low ATP concentrations. As the trailing end of a myosin filament wanders less off axis than at high ATP concentrations, it is less likely to encounter another actin filament and form a crosslink. Therefore, we observed a reduction of long dwell times with short run lengths at low ATP concentrations. Once the myosin filaments reach the plus end of an actin filament, they could contribute to the condensation of actin filaments into clusters by catching and pulling adjacent actin filaments leading to the formation of contractile zones in more crowded regions of the actin filament network (3, 46, 47). The effect of myosin II filaments cross-linking multiple actin filaments on network contractility and entropy production was also reported by the Murrell group (39).

The large dynamic range of iSCAT microscopy allowed us to reveal the dynamics within acto-myosin clusters. Interestingly, myosin II filaments continuously moved into myosin clusters, and within these clusters eventually detached upon reaching the clusters’ center. This resulted in no or only limited growth of clusters over time This would imply that myosin filaments inside clusters are not necessarily in a jammed state and that the turnover of myosin II filaments could contribute to the formation of constitutively active contractile zones. Taken together, our observations of myosin dynamics underline the notion that myosin II filaments can act as motors and crosslinkers, which is important to drive clustering of acto-myosin networks, and that a relatively small number of long-binding, cross-linking myosin II filaments can mediate the transition from the remodeling to the contractile state (Fig. 4C).

## Contributions

DK conceived, designed and performed experiments, purified proteins, analyzed data, wrote manuscript; NH helped in the experiments, analyzed data, wrote manuscript; LM wrote the tracking code and performed analysis; GY and AF built the iSCAT microscopes; MP reviewed manuscript, advised in the development of the tracking code, reviewed/edited manuscript; PK conceived experiments, reviewed/edited manuscript; SM conceived experiments, reviewed/edited manuscript.

## Acknowledgements

DK thanks Madan Rao (NCBS) and Kabir Husain (University of Chicago) for instructive discussions and comments on the manuscript, Mohan Balasubramanian (Warwick Medical School) for material support and instructive discussions, and the Company of Biologists for supporting this work with a Travelling fellowship. NH thanks James Sellers and Yasuharu Takagi from the NIH for providing the actin. DK was funded as a Campus Fellow at NCBS, later as a postdoctoral fellow in the Balasubramanian lab by ERC-2014-ADG N° 671083 and then as an assistant professor in the Wellcome-Warwick Quantitative Biomedical Program by Wellcome Trust grant RMRCB0058. NH was supported by DFG research fellowship, grant HU 2462/1-1, and later by a DFG return grant HU 2462/3-1. LM was supported by Leverhulme Trust Grant RPG-2016-260. GY was supported by a Zvi and Ofra Meitar Magdalen Graduate Scholarship; PK was supported by an ERC Starting Investigator Grant (Nanoscope, 337577). SM and this work were supported by a JC Bose Fellowship from the Department of Science and Technology, and a Margadarshi Fellowship IA/M/15/1/502018 from Department of Biotechnology-Wellcome Trust, India Alliance.

---

**Table 1.** experimental conditions: List of the experimental conditions used in the examples depicted in Figs. 1–4. The table contains the concentration of G-actin, capping protein (CP), myosin II proteins and ATP as well as the average length of actin filaments attached to SLBs, which was measured before the addition of myosin II. Sample size for filament length measurements: N = 505 (Fig.1E-H), N = 151 (Fig 2, 3, 4), N = 255 (Fig S2A, B).

| Table S1. Summary of the experimental conditions |  |  |  |  |  |  |
| --- | --- | --- | --- | --- | --- | --- |
| Figure | [Actin]/ nM | [CP]/ nM | L(F-actin)/ $\mu\text{m}$ | [Myosin II]/ nM | [ATP]<sub>initial</sub>/ $\mu\text{M}$ | $\lambda(\text{laser})$/ nm |
| 1B | 100 | 0 | NA | 0 | 0 | 445 |
| 1C | 0 | 0 | 0 | 20 | 0 | 445 |
| 1E-H | 125 | 0.75 | 3 (+/- 1.4) | 0 | 100 | 445 |
| 2, 3 | 300 | 0 | 7 (+/- 3.5) | 100 | 100 | 635 |
| 4 A, B | 300 | 0 | 7 (+/- 3.5) | 100 | 100 | 635 |
| S2A, B | 250 | 0.5 | 2 (+/- 1.4) | 30 | 100 | 445 |

