## supplementary text for "Myosin II filament dynamics in actin networks revealed with interferometric scattering microscopy"

**Supplementary material**

*Detection of myosin II filaments*

We use two criteria for the detection of a bound myosin filament: particles have to be above an intensity threshold, and they have to display a minimal area characteristic for a single myosin filament oriented along the focal plane. This ensures that the probability to detect a myosin filament that is diffusing freely and not binding to f-actin is very low.

Considering an unbound myosin filament, it would move out of a r = 200nm radius sphere within <10ms, which would not generate enough signal to be considered/ picked up by our tracking algorithm.

This estimate is based on taking the average time for a diffusing particle to reach from the center a point on a sphere with radius r: $\bar{t}= \frac{r^{2}}{6D}$

And with $D= \frac{k_{B}T}{6 \pi\gamma R}$ , where $\gamma=0.89 {10}^{-3} \frac{kg}{s m}$ is the viscosity of water at 23 °C and R = 250nm is the radius of a sphere to represent a myosin filament, we get

$$\bar{t}= \frac{{(0.2 \mu m)}^{2}}{6 (1.48 \frac{{\mu m}^{2}}{s})}=6.7 ms$$

And if we take into account the elongated shape of myosin, with the long semi-axis b = 300nm, a short semi-axis a = 50nm and $D= \frac{k_{B}T\ln(2b/a)}{4 \pi\gamma b}$, we get

$$\bar{t}= \frac{{(0.2 \mu m)}^{2}}{6 (3.31 \frac{{\mu m}^{2}}{s})}=3 ms$$

*Mobility of actin filaments and slippage of myosin II filaments on supported lipid bilayers*

*Here, we are considering the possibility that the motor action of a myosin II filament would lead to a slippage of the actin filament resulting in a reduced net velocity of the myosin filament. For this, we compare the friction of the actin filament created by the actin-membrane linker and compare it to the friction that myosin II filaments experience in aqueous solutions.*

*Similar to considerations described in (30), we can estimate the frictional force on the actin-membrane linker by using the Einstein-Smoluchowski relation, where the drag coefficient is related to its diffusivity D_HKE_  (= 1 µm^2^s^-1^) by:*

$$\gamma_{HKE}= \frac{k_{B}T}{D_{HKE}}$$

*The drag force on a single actin filament could then be estimated as:*

$$F_{actin}= L_{actin}\rho_{HKE}\gamma_{HKE} v_{actin}=L_{actin}\rho_{HKE}\frac{k_{B}T}{D_{HKE}} v_{actin}$$

*If we consider an actin filament of L_actin_=5 µm length, a linear membrane-actin linker density of ρ_HKE_ = 30 µm^-1^ and a maximum velocity of v_actin_ = 400nm s^-1^ (if the myosin would stand still and move actin at maximum speed) the drag force would be*

$$F_{actin}^{max}=0.25 pN$$

*with k_B_= 1.38 10^-23^JK^-1^ and T = 295K.*

*In contrast, the drag force that acts on a myosin filament* *when it moves through an aqueous solution, can be estimated using the drag coefficient of a cylindrical object of length L and radius r moving parallel to the surface in a distance h:*

$$\gamma_{myo}= \frac{2\pi\eta L_{myo}}{ln(\frac{2h}{r_{myo}})}$$

*And the drag force for the myosin filament would be*

$$F_{myo}= \gamma_{myo}v_{myo}= \frac{2\pi\eta L_{myo}}{ln(\frac{2h}{r_{myo}})} v_{myo}$$

*With η= 10^-3^(water), L_myo_ = 550nm, h = 60 nm, r_myo_ = 30 nm, and v_myo_ = 400 nms^-1^ the drag force is*

$$F_{myo}=1fN$$

*That means that the drag on the myosin filament is 250 times smaller than the drag on the actin filament. This explains that myosin filaments move along single actin filaments without causing any significant displacement of actin filaments and that changes in actin filament organization only occur when myofilaments bind to multiple actin filaments resulting to higher friction on myosin II filament motion and force transduction to the actin network.*

***Figure and Video legends***

**Figure S1, Detection and identification of actin and myosin II filaments: A)** Graph illustrating the typical loss of buffer due to evaporation from the experimental chambers at room temperature; data points represent average ± standard deviation of three independent experiments. **B)** Example of median filter to visualize individual actin filaments landing afresh or moving within a static actin network; the median of the image sequence is subtracted from the raw images; 445 nm laser set-up, scale bar: 1 μm. **C)** Histogram of actin filament and myosin II filament contrast measurements with the 635 nm laser iSCAT set-up; actin filaments: N_measure_ = 352 along N_fil_ = 15 filaments and N_myoII_ = 94; black lines depict fit of normal distribution. **D)** Histogram of myosin II filament length measurements with the 635 nm laser iSCAT set-up; black line depicts fit of normal distribution; N = 269. **E)** Image sequence of an actin filament landing on top of another imaged with the 445 nm iSCAT setup. **F)** Corresponding measurement of the interferometric contrast of the region depicted by the white box in (C) showing a stepwise increase of the signal corresponding to the actin filament landing on top of another. **G)** Histogram of the actin layer thickness in a typical experiment based on background-subtracted, local interferometric contrast measurements divided by the average contrast value of an individual actin filament (C) (N=149). **H)** Center of mass trajectory of a 1 µm long actin filament (Fig. 1E, F; Video 3) over time (from dark blue to cyan: 0- 137s), scale bar: 1 μm. **I)** Center of mass trajectory of a 4 µm long actin filament (Fig 1G; Video 4) confined by the surrounding actin network (from dark blue to cyan: 0-200 s), scale bar: 1 μm. **J, K)** Combined plot of MSDs of actin filaments shorter than 1.5µm (**J**) or longer (**K**).

**Figure S2, Myosin II filament dynamics: A, B)** Image sequence depicting the dynamics of a membrane bound acto-myosin network at (**A**) the start of the experiment, in the remodeling state at 100 μM ATP and (**B**) after 16min, when the network becomes contractile; images were acquired with the 445 nm laser iSCAT setup; scale bar: 2 μm. **C)** Image sequence showing the run of two myosin II filaments (black) on actin filaments; 635nm laser iSCAT setup, scale bar: 1 μm. **D)** Corresponding kymograph obtained along the red lines in (C), black regions correspond to myosin II filaments that are moving along actin filaments; time color bar: 36 s, scale bar: 1 μm. **E)** Plot of run length vs binding time of myosin II filaments at t = 1 min (100 μM ATP) (Fig. 2E). **F)** Plot of run length vs binding time of myosin II filaments at t = 16 min (Fig. 2H). **G)** Boxplot depicting the velocity of actin filaments in single regions of the acto-myosin network indicating a very limited impact of actin filament displacement on the myosin II filament velocity measurements.

**Figure S3, Myosin II filament single particle tracking: A-C)** Example of the routine to classify track segments of directed or random motion: based on the tracking data, the angular difference between particle orientation and direction of motion (**A**), the straight-line distance from the particle’s origin (**B**) and the average correlation in displacements between adjacent timesteps (forward: $C_{+}=\frac{\boldsymbol{dr}_{t+2\Delta t}\cdot\boldsymbol{dr}_{t+ \Delta t}+ \boldsymbol{dr}_{t+\Delta t}\cdot\boldsymbol{dr}_{t}}{2}$, backward: $C_{-}=\frac{\boldsymbol{dr}_{t-2\Delta t}\cdot\boldsymbol{dr}_{t- \Delta t}+ \boldsymbol{dr}_{t-\Delta t}\cdot\boldsymbol{dr}_{t}}{2}$) were calculated (**C**) to reduce false positive detection (22); segments of correlation of displacement above the threshold (here 0.5 nm^2^) were classified as directed motion (blue dotted line), whereas segments below the threshold were deemed to be of random motion (red dotted line ). **D)** Plot of the average (±std) number of automatically detected myosin II filaments per frame in the recordings at the indicated timepoints (N > 250 frames for each condition). **E)** Plot of the ratio of the numbers of detected myosin II filaments that exhibit each binding time at t=16 min (contractile state) and t = 1 min (remodeling state), indicating a 1.6-fold increase for binding times longer than 3 s; counts were normalized to the total number of detections at t = 1 min and t = 16 min. The strong fluctuations at times longer than 10s are due to the small numbers (< 10) of detected events. **F)** Box plot depicting the ratio of the duration of directed motion versus dwell time for each myosin II filament displaying directed motion at the indicated time points (N(1, 5, 9, 13, 16 min) = [221, 374, 158, 608, 640]). **G)** Histogram of the switches of each tracked myosin between directed and undirected motion at the indicated time points (sample numbers are similar as in F). **H)** Displacement per frame distribution of myosin II filaments in directed motion at the indicated time points of the experiment. **I)** Radial plot depicting the angular difference α between myosin II filament orientation and its velocity vector for all detected myosin II filament steps at the indicated time points. **J)** Radial histogram plot depicting the ratio of steps at directed motion versus all detected steps for each angular difference α at indicated time points. **K)** Plot of the average (±std) effective torsional spring constant of myosin II filaments at the indicated time points; std estimation is based on the error of myosin orientation detection.

**Video 1** Movie recorded with a 445nm laser iSCAT system showing the formation of an SLB by landing and fusion of single uni-lamellar vesicles on a glass slide. The reduced contrast at the end of the video indicates the presence of the SLB as it smoothens the glass roughness.

**Video 2** Movie showing the increase in interferometric scattering when an actin filament lands on top of another (Fig. S1E, F)

**Video 3** Movie showing diffusion of a 1 µm long actin filament within a membrane-bound actin network (removed by median filtering) (Fig. 1E, F and Fig. S1H)

**Video 4** Movie showing diffusion of a 4 µm long actin filament within a membrane-bound actin network (removed by median filtering) (Fig. 1G and Fig. S1I)

**Video 5** Movie showing the transition of the acto-myosin network from a remodeling to a contractile state upon depletion of ATP over time (Fig. S2A, B).

**Video 6** Movie showing the release of myosin II filaments from actin after addition of fresh ATP (100 µM) indicating that the contractile state of myosin II filaments in this set up is reversible.

**Video 7** Movie showing myosin II filament motion along actin filaments (Fig. S2C, D). Scale bar 1 µm.

**Video 8** Movie showing tracking of a single myosin filament (Fig. 3A, B; S3 A-C). Scale bar 1 µm.

**Video 9** Movie showing the flow of myosin II filaments into and within an acto-myosin cluster after subtraction of the time median (Fig. 4 A, B). Scale bar 2 µm.
