## Supplementary figures and images for "Myosin II filament dynamics in actin networks revealed with interferometric scattering microscopy"

### supplementary figures

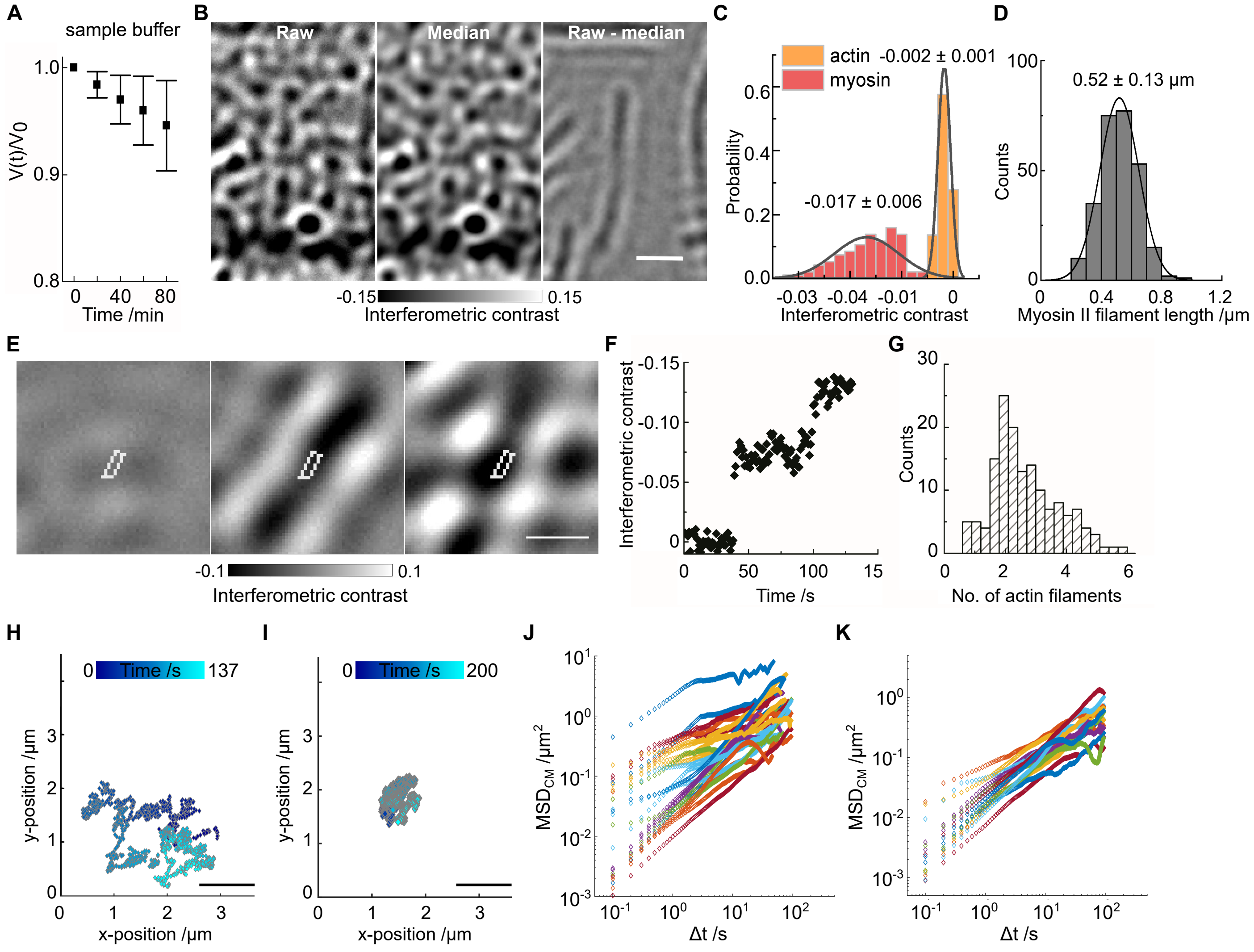

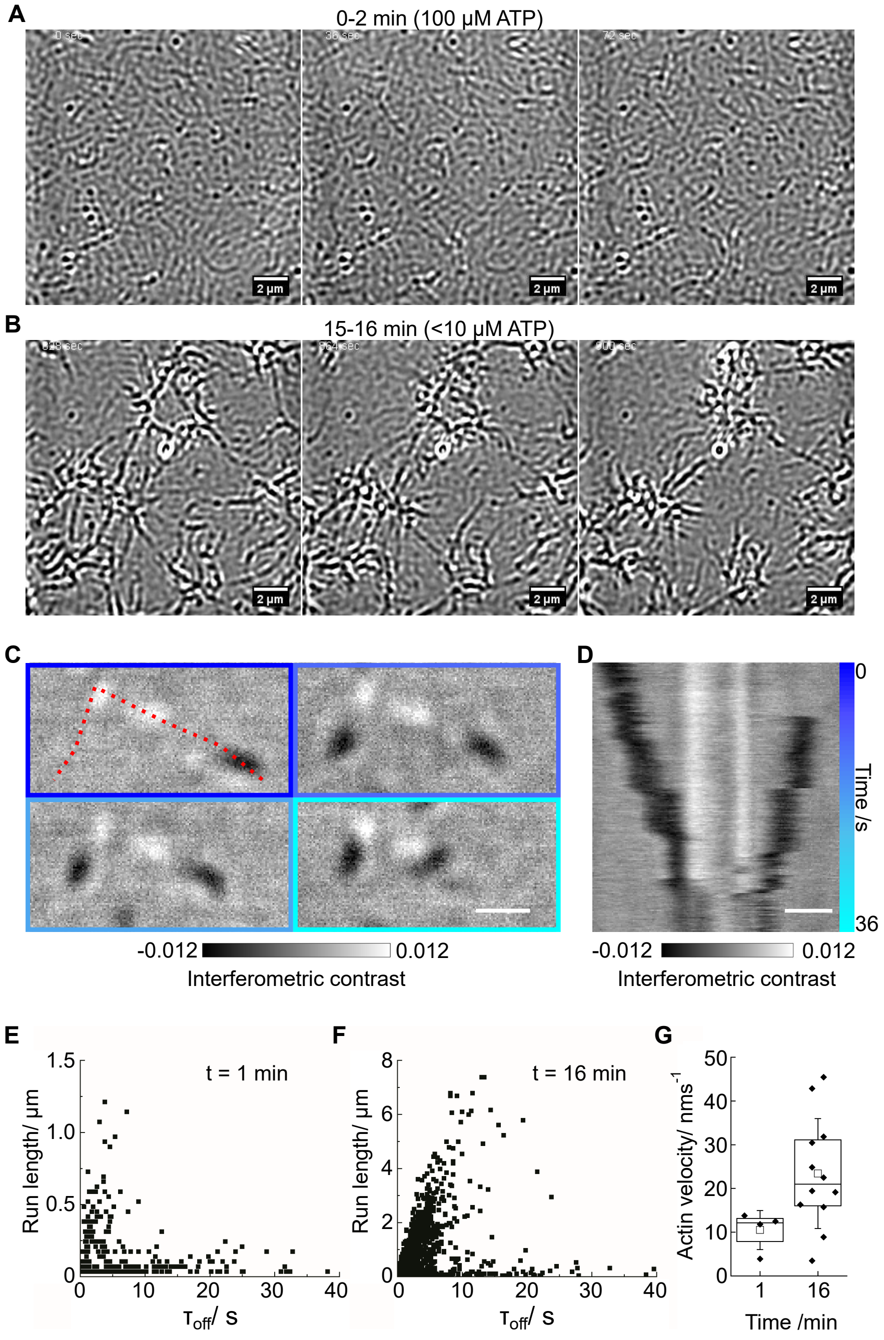

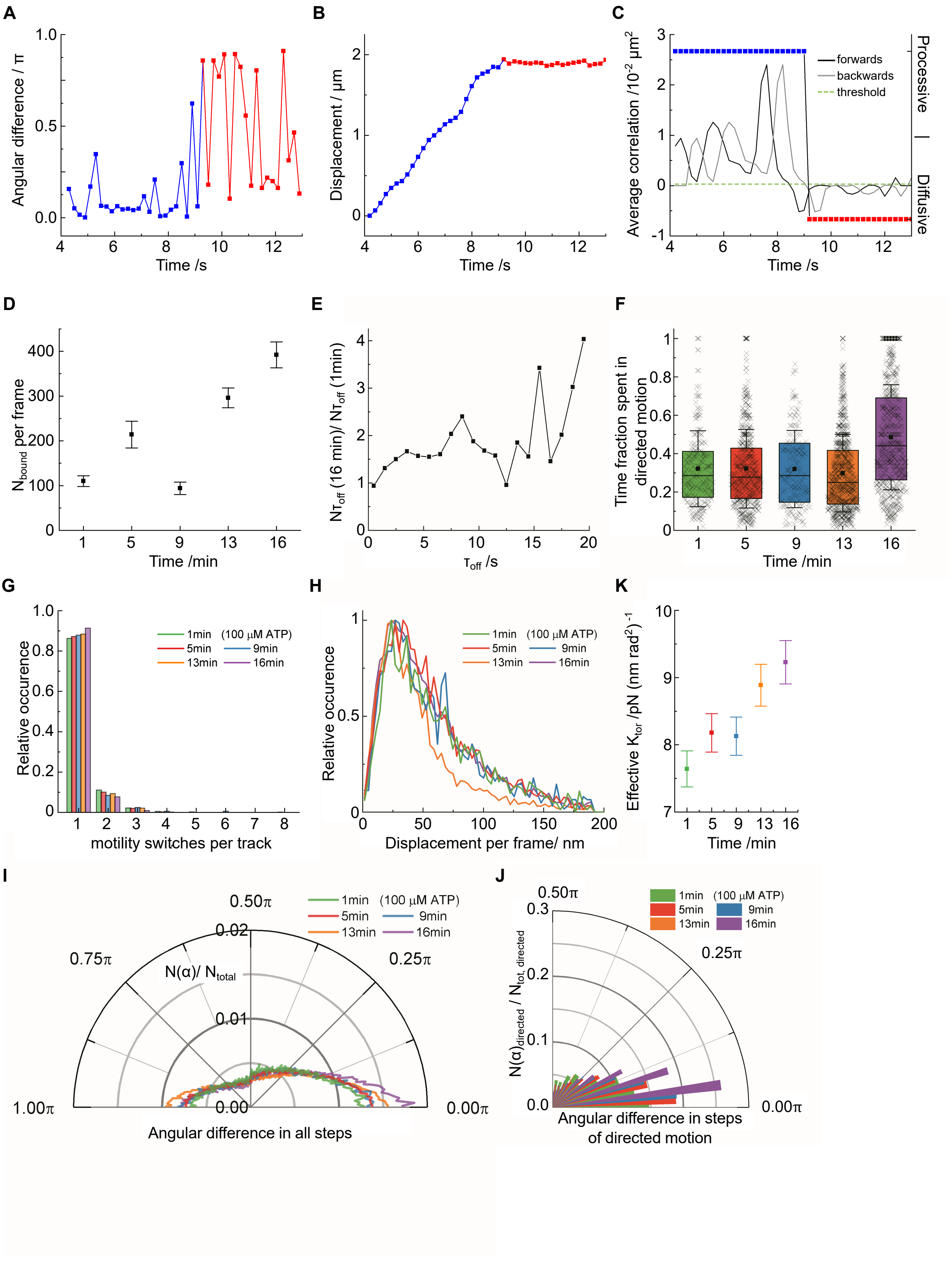
